# A SNP affects Wnt4 expression in endometrial stroma, with antagonistic implications for pregnancy, endometriosis and reproductive cancers

**DOI:** 10.1101/2022.10.25.513653

**Authors:** Mihaela Pavlicev, Caitlin E. McDonough-Goldstein, Andreja Moset Zupan, Lisa Muglia, Yueh-Chiang Hu, Fansheng Kong, Nagendra Monangi, Gülay Dagdas, Nina Zupancic, Jamie Marziaz, Debora Sinner, Ge Zhang, Günter Wagner, Louis Muglia

## Abstract

The common human single nucleotide polymorphism rs3820282 is associated with multiple phenotypes ranging from gestational length to likelihood of endometriosis and ovarian cancer and can thus serve as a paradigm for a highly pleiotropic genetic variant. Pleiotropy makes it challenging to assign specific causal roles to particular genetic variants. Deleterious mutations in multifunctional genes may cause either the co-occurrence of multiple disorders in the same individuals (i.e., syndromes), or be repeatedly associated with a variety of disorders in a population. Moreover, the adverse effects can occur in combination with advantages in other traits, maintaining high frequencies of deleterious alleles in the population. To reveal the causal role of this specific SNP, we investigated the molecular mechanisms affected by rs3820282 in mice. We have shown previously that rs3820282 introduces a high affinity estrogen receptor 1 binding site at the *Wnt4* locus. Having introduced this nucleotide substitution into the homologous site of the mouse genome by CRISPR/Cas 9 we show that this change causes a specific upregulation of *Wnt4* transcription in the endometrial stromal cells during the preovulatory estrogen peak in late proestrus. Transcriptomic analysis of the whole uterus reveals broad systemic effects on uterine gene expression, including downregulation of proliferation and induction of many progesterone-regulated pro-implantation genes. The effect on proliferation is limited to the luminal epithelium, whereas other effects involve the uterine stromal compartment. We suggest that in the uterus, these changes could contribute to increased permissiveness to embryo invasion. Yet in other estrogen-responsive tissues, the same changes potentially lead to decreased resistance to invasion by cancer cells and endometriotic foci. A single molecular effect of rs3820282 on *Wnt4* expression may thus underlie the various associated phenotypic effects.

## Introduction

Genome-wide association studies (GWAS) frequently reveal associations between multiple diseases and a polymorphism at the single genomic locus, suggesting pleiotropy ^1–4^. Disease pleiotropy may originate from either the same molecular mechanism involved in different disease processes or through broader systemic effects of the mutation - such as immune or metabolic disorders - affecting different disease phenotypes. Alternatively, different genetic variants involved in etiologically unconnected diseases could be co-inherited due to genetic linkage ^3^. Beyond the mere presence of genetic correlation, recent studies have also addressed the *direction* of the effects of specific variants on single traits (diseases), i.e., whether they are deleterious or beneficial with respect to different phenotypes. While many pleiotropic variants have congruent fitness effects in all diseases (i.e., the same allele is deleterious in all), studies also commonly find cases of antagonistic pleiotropy^5^, where a variant augments the likelihood of one disorder, while protecting the carrier from another ^6–9^. Antagonism in pleiotropy is particularly interesting as it may explain the high frequency of some of the apparently deleterious alleles in a population^10^, reflect a biological trade-off that can inform the disease mechanisms, and provide guidance in search of treatment strategies that should be preferred or avoided due to the associated side effects ^3^.

Deciphering the molecular basis of statistical associations requires determining the exact genomic location of the causal polymorphism, the target gene (most variants are in the regulatory genome), the spatial and temporal context of the mutational effect (i.e., tissue or cell type, stage of development or physiology), the molecular pathway mediating the effect, and the organismal processes mapping the genetic change onto the final phenotype. Information from multiple diseases can facilitate the analysis of the mechanistic basis of mutation effects^11^. In this paper, we present the results of a functional genetic study of one single nucleotide polymorphism (SNP) located in the first intron of *Wnt4* (rs3820282) on human chromosome 1 (1p 36.12). The frequency of the alternate allele at rs3820282 (T, with reference allele being C) varies strongly across human populations, ranging from less than 1% in Africa to over 50% in SE Asia^12^. This SNP has been associated with variation in multiple human female reproductive traits. Thereby, the same allele is associated with deleterious as well as protective effects on diseases such as endometriosis^13–16^, breast cancer^17^, ovarian epithelial cancer^18^, length of gestation^19^, and leiomyoma^20–23^. Specifically, the alternate allele has been associated with higher likelihood of endometriosis, fibroids (leiomyoma), and ovarian epithelial cancer, yet it is also associated with longer gestation and is potentially protective against preterm birth. Rather than focusing on the etiology of any one disease, we focus here on the immediate molecular and cellular mechanisms through which the locus exerts its function, likely mediating different disease contexts.

The disease associations of this SNP are consistently related to female reproductive biology, a context in which regulation by estrogen is central. An exceptionally well powered GWAS, in an attempt to understand the genetic basis of preterm birth, found that the alternate allele introduces a potent binding site for estrogen receptor alpha (ESR1)^19^. Moreover, this binding site overlaps with open chromatin in immortalized human endometrial stromal cell line, HESC^24^, in vitro^19^. The information on downstream consequences of mutation is less robust, although this question has been experimentally addressed previously (e.g.,^17,25^, see discussion).

Estrogen responsiveness has been directly implicated in most diseases associated with the genomic region of the rs3820282 allele. Endometriosis has been associated with the incomplete transition of the endometrium from the estrogen-dominated proliferative phase to the progesterone-dominated secretory phase of the estrous cycle, also referred to as progesterone resistance^26^, due to dysregulation in the early secretory phase of many genes mediating progesterone effects. Fine-mapping studies of the region to identify the causal SNP in the context of endometriosis have confirmed rs3820282 as a prioritized candidate^16,27^. Estrogen also enhances the growth of uterine leiomyoma (fibroids), a common benign tumor of the myometrial smooth muscle^28^. A recent GWAS in a large Icelandic cohort found associations of uterine fibroids with alleles at two sets of loci: loci shared with a wide range of cancers, and loci shared with various reproductive abnormalities, specifically implicating involvement of estrogen signaling, among them rs3820282^23^. Increased estrogen and androgen, as well as hyperactivity of stroma have further been proposed to contribute to the pathogenicity of ovarian cancer^29^ and recent association studies linked both ovarian as well as breast cancers to the region of the focal locus^23^.

This genomic region of interest is conserved across placental mammals (Fig. 1), likely due to the key roles of Wnt in vertebrate development and physiology^30^. The mammalian reference genomes (including C57B/6J mouse line, used in this study as wild type, WT) carry what in the human population is the major allele. Many aspects of female reproductive biology are also conserved between mice and humans. Therefore, we expect that the molecular effect of this mutation in mice will be informative for studying its effect in humans.

**Figure 1.**
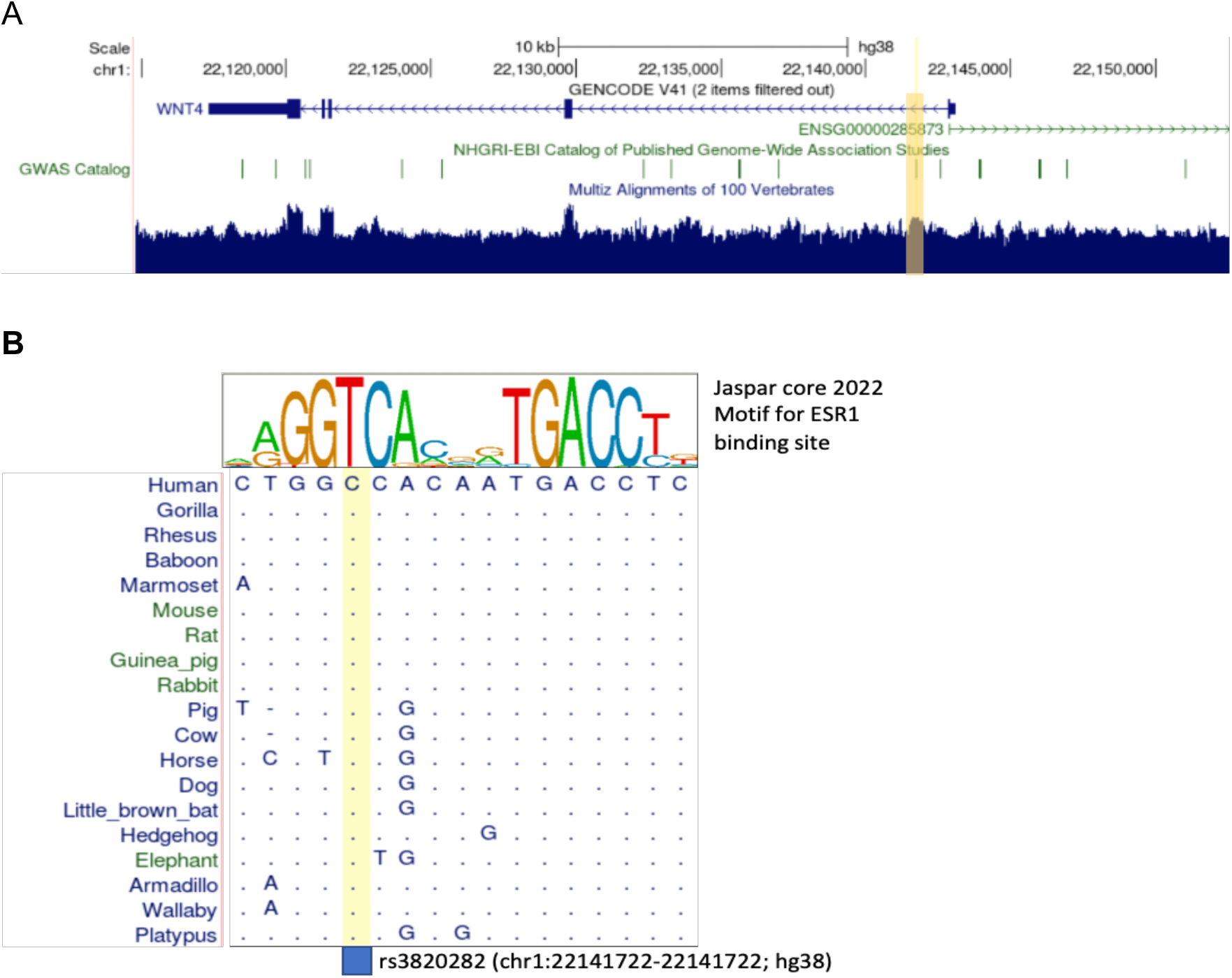
Alignment of the noncoding region around rs3820282 across mammalian genomes (UCSC genome browser). A) overall conservation of the *Wnt4* region. B) The binding site motif for ESR1 on top summarizes the sequence variation across the functional binding sites.The position of rs3820282 is shown in yellow in A and B, and changes the human nucleotide from C to T.

To determine the molecular mechanisms affected by the SNP rs3820282, we generated CRISPR/Cas9-modified transgenic (TG) mouse line, homozygous for what is the human alternate allele, and compared it to the homozygous WT mouse line of the same background. Genome-editing circumvents the effect of linkage between the neighboring SNPs, due to which causal mutations are hard to distinguish from the linked ones in the human population. This method aids in the feasibility of experimentation and allows us to avoid the heterogeneity of the genetic background between individuals, which can hinder the attribution of effects to any one polymorphism. The context-dependent effects of a polymorphism is a relevant aspect that must be addressed separately. Using this transgenic mouse line we reveal that the mutation enhances the periodic estrogen-induced uterine preparation for implantation. We propose that this effect, when appearing in other estrogen-responsive tissues, may similarly enhance the permissibility of tissues to endometriosis and metastasis.

## Results

We used gene editing to replace the single nucleotide mouse wild type allele with the human alternate allele at the location in the mouse genome corresponding to rs3820282 in humans, using CRISPR/Cas9 (see Methods for detail). The position rs3820282 and its flanking sequences are 98% conserved between the human and mouse reference genomes (Fig. 1). Live born pups were genotyped by PCR and then further confirmed by Sanger sequencing. Two out of three lines in which the overall region had been modified, had the specific one-nucleotide substitution we aimed for. While we focus on only one of the two genome-edited mouse lines in this work for the full range of analyses, both show the same relevant phenotype described below: the effect on Wnt4 expression in proestrus (Fig. 1, Fig. S1). We note that we did not observe differences in gestation length (WT: 19.3 ± 0.06 days, N = 7, KI-1: 19.2 ± 0.07 days, N = 5, KI-2: 19.2 ± 0.04 days, N = 7, P = 0.33; Wilcoxon-Mann-Whitney rank sum test) or litter size (WT: 8.4 ± 0.65 pups, N = 7, KI-1: 7.8 ± 0.58 pups, N = 5, KI-2: 8.2 ± 0.56 days, N = 7, P = 0.66) in either transgenic line (Fig. S2).

### Transgenic allele at the rs3820282 upregulates uterine Wnt4 mRNA expression in late proestrus

Given the previous finding that rs3820282 SNP is enhancing ESR1 binding^19^, we investigated which genes may be regulated by this novel binding site. We focused on the uterine tissue, as previously reported associated reproductive phenotypes involved the uterus. We first investigated the expression of two genes adjacent to the SNP, *Cdc42* and *Wnt4*. We analyzed how the SNP affected the uterus across multiple stages of the ovarian cycle and pregnancy, specifically proestrus, estrus, and early pregnancy (7.5 days post conception, dpc). The mouse uterus undergoes important changes in preparation for implantation even prior to mating, thus pregnancy success can be influenced both during the pregnancy, as well as pre-pregnancy, during the stages preceding fertilization and implantation. We found no effect on the *Cdc42* gene adjacent to the polymorphism (proestrus: N = (11 TG, 4 WT), P = 0.34; estrus: N = (6 TG, 2 WT); Wilcoxon-Mann-Whitney rank sum test, Fig. 2).On the other hand, the two genotypes differed only in the uterine expression of *Wnt4* during proestrus, with a 1.48 log2 fold increase in *Wnt4* expression in the transgenic compared to the wild type (N = (14 TG, 8 WT), P = 0.002). No significant difference between the genotypes was observed in Wnt4 mRNA expression in either estrus (N = (6 TG, 5 WT), P = 0.08, Fig. 2) or early pregnancy (7.5 dpc, N = (3 TG, 3 WT), P = 0.70, Fig. S3). Differences in *Wnt4* expression were also not observed in the ovary at either proestrus (N = (3 TG, 5 WT), P = 0.57, Fig. S4) or estrus (N = (5 TG, 3 WT), P = 0.79, Fig. S4)

**Figure 2.**
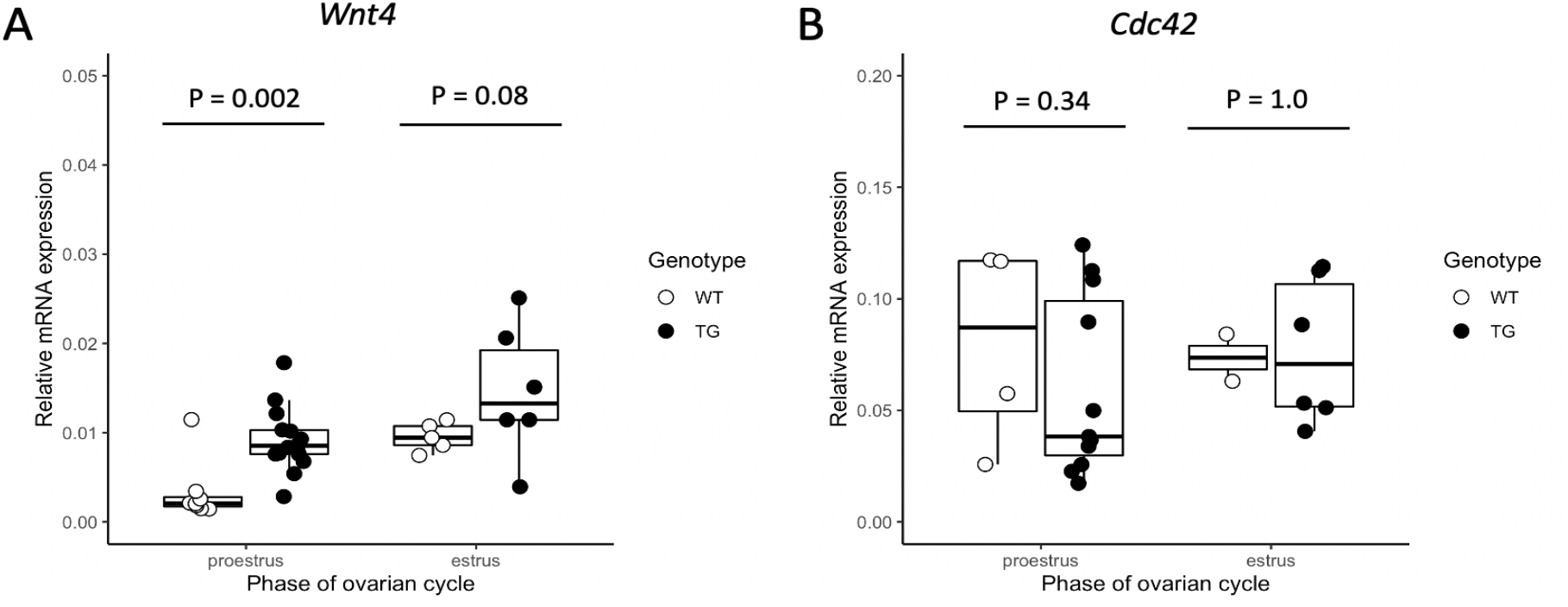
rs3820282 transgenic SNP influences expression of *Wnt4* in the uterus. (A) The mouse model of the rs3820282 transgenic (TG) SNP had significantly higher levels of *Wnt4* in the proestrus phase of the ovarian cycle (p = 0.002) compared to the wildtype (WT). Although differences in expression were not significant in the estrus phase (p = 0.08). (B) Other adjacent genes to the SNP such as *CDC42* did not differ between genotypes (p > 0.2).

### Endometrial stromal fibroblast is the cell type affected by the transgenic allele

To determine the uterine cell type in which *Wnt4* is upregulated in the transgenic line, we performed *in situ* hybridization using RNAscope^31^ on proestrus uteri. We found that *Wnt4* is expressed in luminal and glandular epithelium in both genotypes. In contrast, in the transgenic line *Wnt4* is also strongly expressed in the stromal cells specifically underneath the luminal epithelium (Fig. 3A). To further solidify this finding, we isolated primary endometrial stromal fibroblasts during late proestrus from transgenic and wild type mice and measured the respective expression levels of *Wnt4* by qPCR. Primary cells isolated from the transgenic line exhibited a 2.71 log2 fold upregulation of *Wnt4*, relative to those of a wild type (N = (5 TG, 6 WT), P = .0004; Fig. 3B). Stronger upregulation of *Wnt4* in the primary endometrial stromal cell culture relative to the bulk uterine transcriptome is consistent with the former being enriched for the affected cell type.

**Figure 3.**
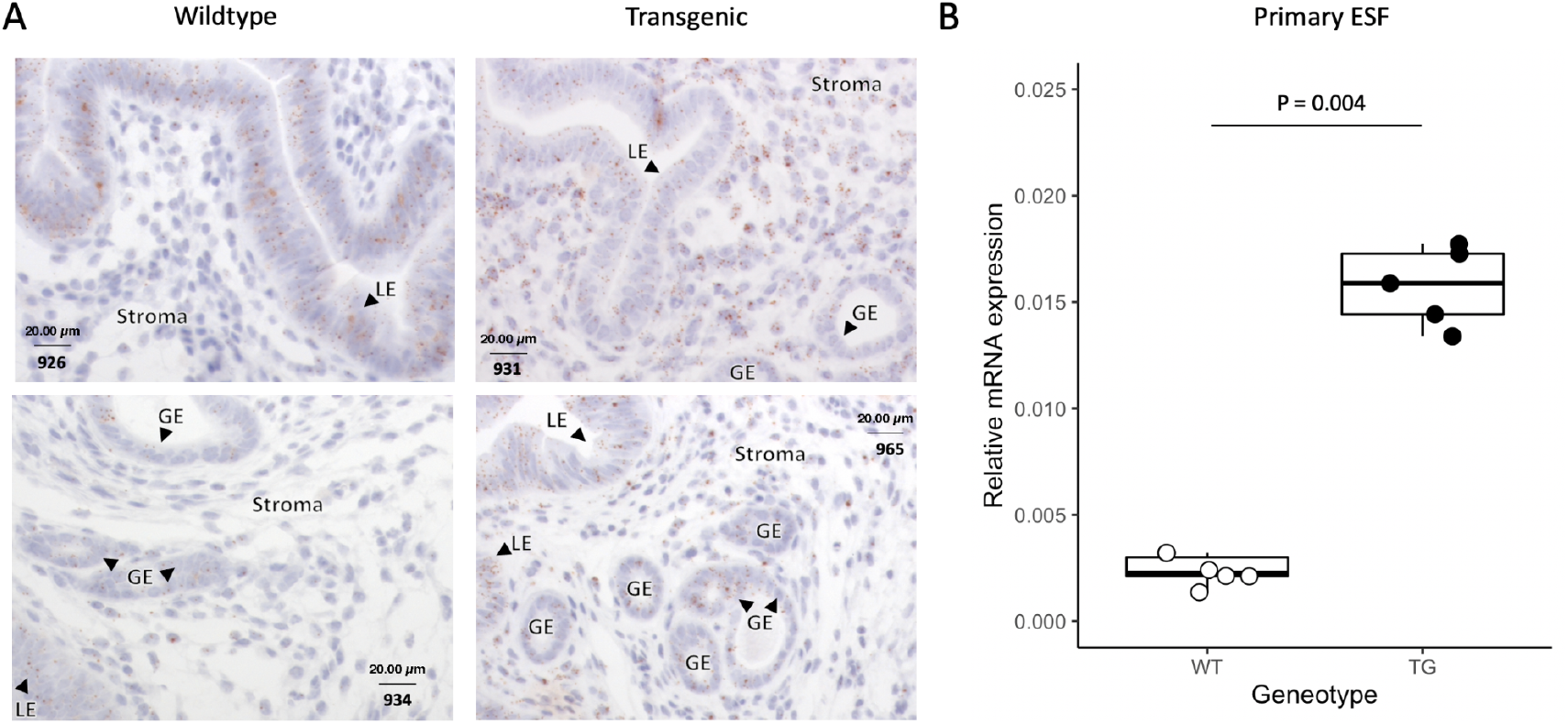
Changes in *Wnt4* proestrus expression are concentrated in the uterine stroma. A) Using RNAscope we localized the expression of *Wnt4* to the luminal epithelium of both wildtype in transgenic lines during proestrus. Expression was also identified in the stroma for the transgenic line but was not detectable for the wildtype. Luminal epithelia (LE), glandular epithelia (GE), and stroma are indicated. Two biological replicates are shown for each line. Wnt4 is stained a light brown and the nucleus is a light indigo, magnification at 40x. B) The concentration of *Wnt4* expression differences in the stroma were confirmed with qPCR of primary endometrial stromal fibroblast cells.

### Transcriptomic differences suggest the mechanisms affected by the mutation

To understand the overall uterine consequences of the mutation, we compared the uterine tissue-transcriptomes of the WT and the homozygous TG mice in proestrus, the only stage which showed changed *Wnt4* expression. This approach revealed the systemic modification of the uterine transcriptome due to increased *Wnt4* in the individuals carrying the transgenic alleles (Fig. S5).

Uterine transcriptomes showed upregulated *Wnt4* expression in transgenic individuals (1.8-log2 fold higher expression in the TG strain, adj. P < 0.001), congruent with qPCR results. We detected 142 genes to have significantly changed proestrus uterine expression levels in the TG relative to WT line (adj. P < 0.05; Table S1 in Supplementary material; Fig. 4A). Of these, five times more genes are upregulated (119) than downregulated (23), a significant enrichment for increased expression (P < 0.001, exact binomial test; Fig. 4B). In the following, we focus on the major GO groups of genes with changed expression, and their biological roles in the uterus. Their pathogenic potential will be addressed in the discussion.

**Figure 4.**
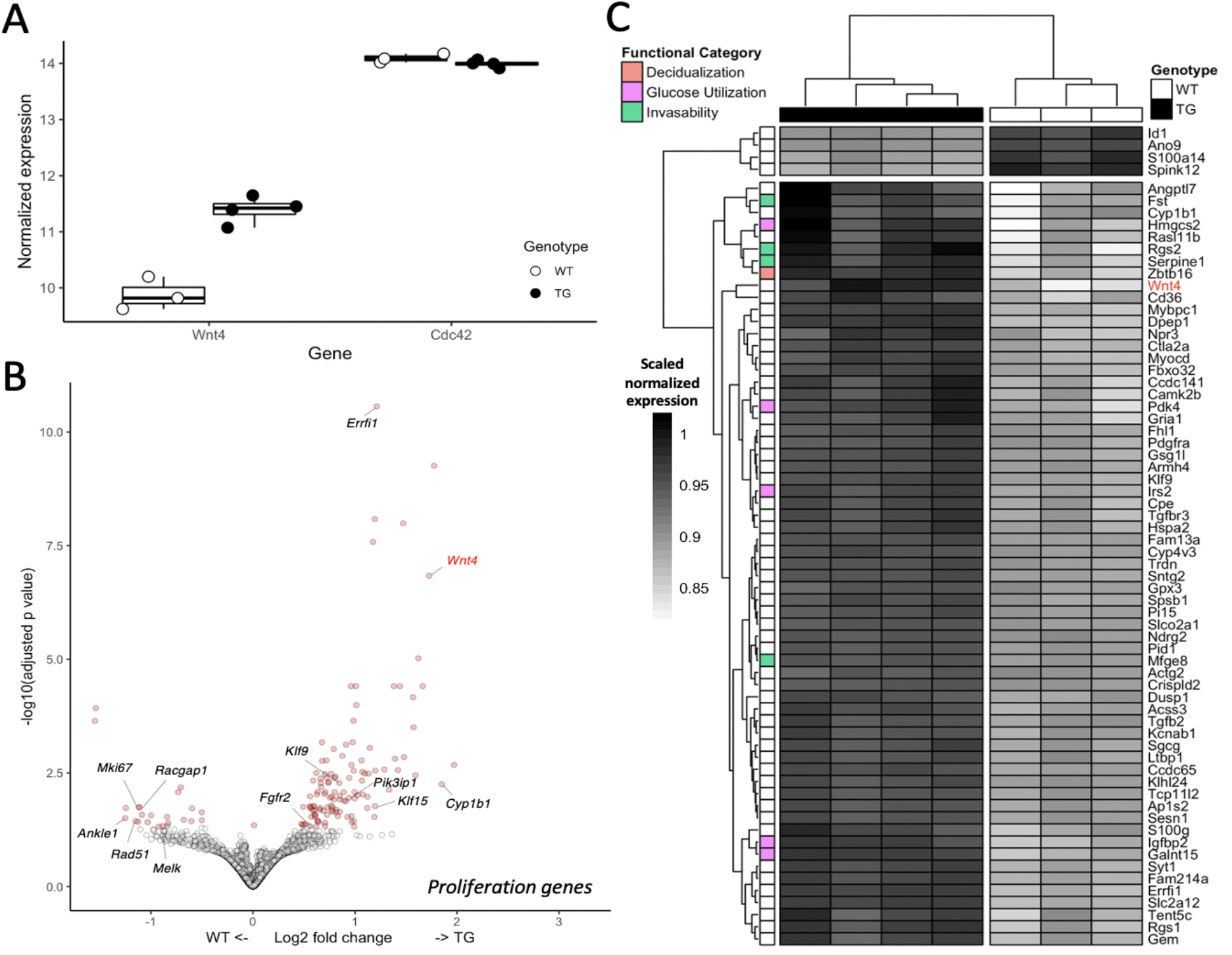
SNP mediated differential expression in the transgenic proestrus uterus. A) Effect of the SNP on flanking genes, with differences in *Wnt4* expression between the wildtype and transgenic lines, but not *Cdc42*, were confirmed in the RNAseq analysis. B) Volcano plot showing significance and effect size distribution. Significantly differentially expressed genes (adj. P > 0.05) are in red. Wnt4 is indicated in red and genes associated with proliferation are labeled. Genes that are associated with increased proliferation were found to decrease in expression (*Mki67, Racgap1, Ankle1, Rad51,* and *Melk*) whereas genes that are associated with the progesterone mediated repression of proliferation increased in expression (*Errfi1, Klf9, Pik3ip1, Klf15, Fgfr2,* and *Cyp1b*1). C) Heatmap of significantly differentially expressed genes with an absolute log2 fold change > 0.5. Genes with functions of relevance to reproductive phenotypes were among those with the greatest increase in expression. In particular, invasibility (*Fst, Rgs2, Serpine1, and Mfge8*), decidualization (*Zbtb16)*, and glucose utilization (*Hmgcs2, Pdk4, Irs2, Igfbp2,* and *Galnt15*).

#### Downregulated genes are enriched from proliferation pathways

The significantly downregulated genes showed very clear enrichment in genes related to proliferation, mitosis and DNA damage repair (e.g., *Mki67, Racgap1, Tacc3, Ankle1, Rad51, Melk;* Fig. 4B). Specifically, hallmark gene set enrichment^32^ was found for processes involving the cell cycle including the G2/M checkpoint, DNA damage repair, and genes associated with mitotic spindle assembly (adjusted p < 0.001; Fig. S6; Table S2). In addition, upstream proliferation effectors were among hallmark gene set enrichments in downregulated genes, including targets of proliferation-inducing E2F and Myc transcription factors, unfolded protein (a cellular ER stress response), and Mtorc1 complex. Immunohistochemistry with anti-MKI67 (proliferation marker) antibody showed that the protein expression of MKI67 is limited to the luminal epithelium in both genotypes, and hence the downregulation of proliferation too must be limited to the epithelium (Fig. 5). Relevant to the disease phenotypes associated with the SNP, there is a significant downregulation of *Spink12*, the serine protease inhibitor, known to be expressed in various epithelia and regulated by differentiation signals. A related family member *Spink13* was shown to be protective against ovarian cancer invasion^33^. Also downregulated is *Akr1c19* (mouse ortholog of human *AKR1C1*), the progesterone metabolizing enzyme.

**Figure 5.**
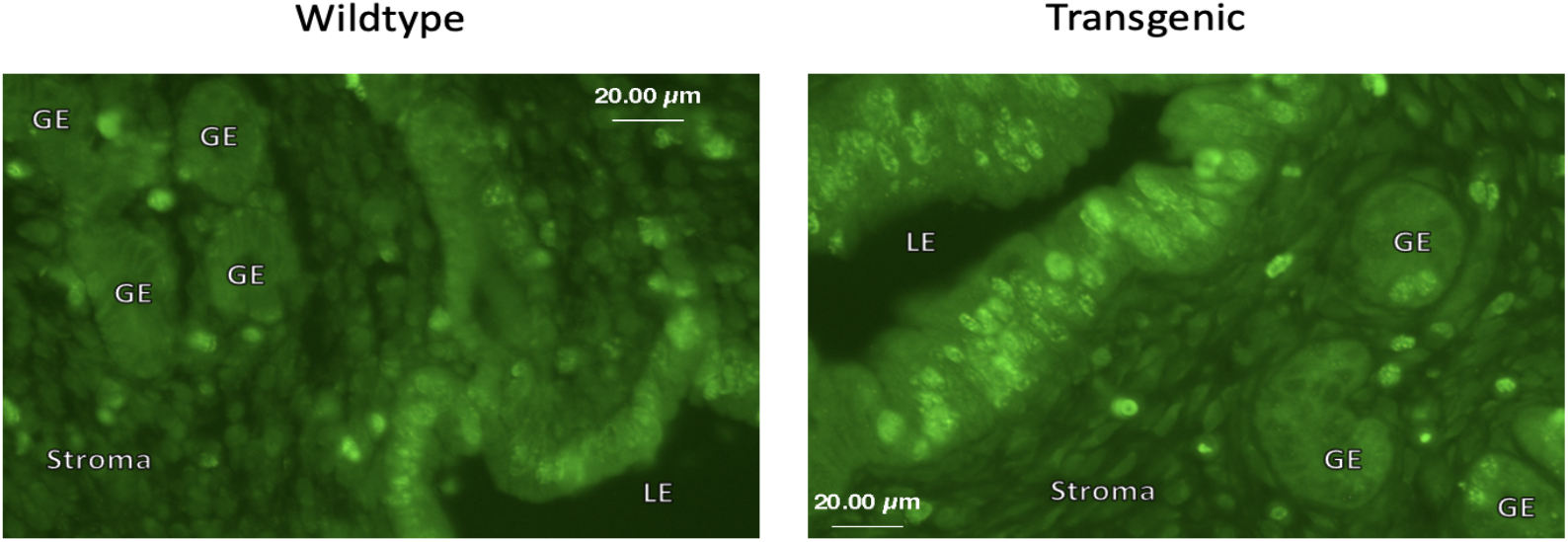
Proliferation marker expressed primarily in the proestrus luminal epithelium. Marker of proliferation, MKI67 is expressed in both genotypes in the proestrus luminal epithelium. Thus, the influence of decreased *Mki67* gene expression in the transgenic line is most likely restricted to the epithelium. Magnification at 40x.

#### Upregulated genes are enriched for processes involved in stromal decidualization, invasibility, suppression of epithelial proliferation and aspects of the progesterone response

Upregulated genes indicate substantially more complex effects of increased *Wnt4* expression than the downregulated genes (Fig. 4C). The enriched hallmark gene sets predict upregulation of pathways related to epithelial-mesenchymal transition, myogenesis, hypoxia, coagulation, KRAS-signaling and TNFA signaling via NFKB, among others (adj. P < 0.001; Fig. S6; Table S2). These pathways are involved in normal uterine processes, with upregulated genes shown to be crucial for implantation including: decidualization (*Zbtb16, Tcf23, Crebrf*), invasibility of uterus by the trophoblast (*Fst, Rgs2,* Integrins), and hemostasis/vascularization *(Rgs2, F5, Serpine1, Mfge8*). In the following we briefly address the reported functions of the most upregulated genes.

We observe in the transgenic line the upregulation of many genes previously associated with progesterone increase, such as occurs in both mouse (early pregnancy) and human (early secretory phase), but without observing upregulation of progesterone receptor. In particular, we identify several genes related to modulation of uterine epithelial proliferation, a consequence of progesterone increase (Fig. 4B). The most significantly upregulated factor involved is *Errfi1*, the Errb feedback inhibitor 1. Errfi1 has been shown to inhibit the action of epidermal growth factors. Accordingly, the ablation of *Errfi1* leads to uterine epithelial hyperplasia^34^. In the uterus, *Errfi1* was shown to mediate progesterone activity by inhibiting the mitogenic action of Erbb2 (receptor tyrosine kinase) and opposing the estrogen-driven endometrial proliferation in the presence of progesterone^35^. Further upregulated genes, such as Krüppel-like factors *Klf15* and, to a lesser extent, *Klf9* mediate suppression of epithelial growth upon combined estrogen and progesterone treatment, enhancing the pro-implantation actions of progesterone^36^. Knockout of *Klf9* inhibits implantation, thus we reason that its upregulation may positively affect implantation^37–39^. Similarly, *Pik3ip1* inhibits Pki3-mediated activation of proliferation and is progesterone regulated^40,41^. Fgfr2 is another upregulated negative regulator of epithelial proliferation^42^. The enhancement of multiple proliferation inhibitors is consistent with the downregulation of proliferation-enhancing genes and the decrease in proliferation marker *Mki67*, whose expression is limited to epithelium. Consistent with progesterone-response-like expression pattern is also the upregulation in transgenic animals of *Cyp1b1*, an enzyme metabolizing estrogen. This may contribute to the local metabolism of mitogenic estrogen mediating the antagonistic effects of progesterone and estrogen on proliferation.

Several upregulated genes are involved in decidualization, a uterine endometrial transformation that in mice does not normally precede fertilization (but does in humans). Among these is *Zbtb16*, a transcriptional repressor involved in cell cycle control, and shown to mediate progesterone receptor-driven decidualization of human endometrial stromal cells^43^. Also upregulated are *Tcf23*, a decidualization-associated transcription factor^44^, and *Crebrf*, a factor involved in regulation of stress response, whose expression is known to increase at implantation site^45^.

Among the significantly upregulated genes known for their roles in modulating receptivity and invasibility is Folistatin (*Fst*), whose uterine-specific knockout in mouse reduces the responsiveness of the luminal epithelium to estrogen and progesterone^46^. Similarly, integrins (*Itgb3, Itga8,* and associated *Lim2*) have been repeatedly implicated in embryo attachment^47,48^, and are also associated with wound healing and vascularization^49^, the epithelial-to-mesenchymal transition (EMT) and stromal invasibility in cancer^50^. Also upregulated in the transgenic mice are the regulators of G-protein signaling, *Rgs1* and *Rgs2*. These proteins have been linked to regulation of vascular tone in implantation. Rgs2 increases at the implantation sites as stromal cells start to decidualize in mouse, and its downregulation inhibits trophoblast growth *in vitro^51^*. Further indication of early vaso-activation is the upregulation of genes with endothelial or peri-endothelial expression (*Serpine1^52^, Mfge8^53^*).

Notable as well is the increased expression of genes involved in glucose utilization (*Irs2, Igfbp2, Pdk4, Hmgcs2, Slc2A12;* Fig. 4C). Aberrant glucose metabolism has been associated with defects of decidualization and implantation, likely because glucose contributes to protein glycosylation of abundant glycoproteins in the uterus, including those of glandular secretions and also serves as critical nutritional support for developing embryo. Indeed, we found that the transgenic uteri appear to produce higher amounts of polysaccharide-rich glandular secretions (Fig. 6). In accordance with this observation, we note upregulation of *Galnt15*, an enzyme catalyzing protein glycosylation, in particular of mucin.

**Figure 6.**
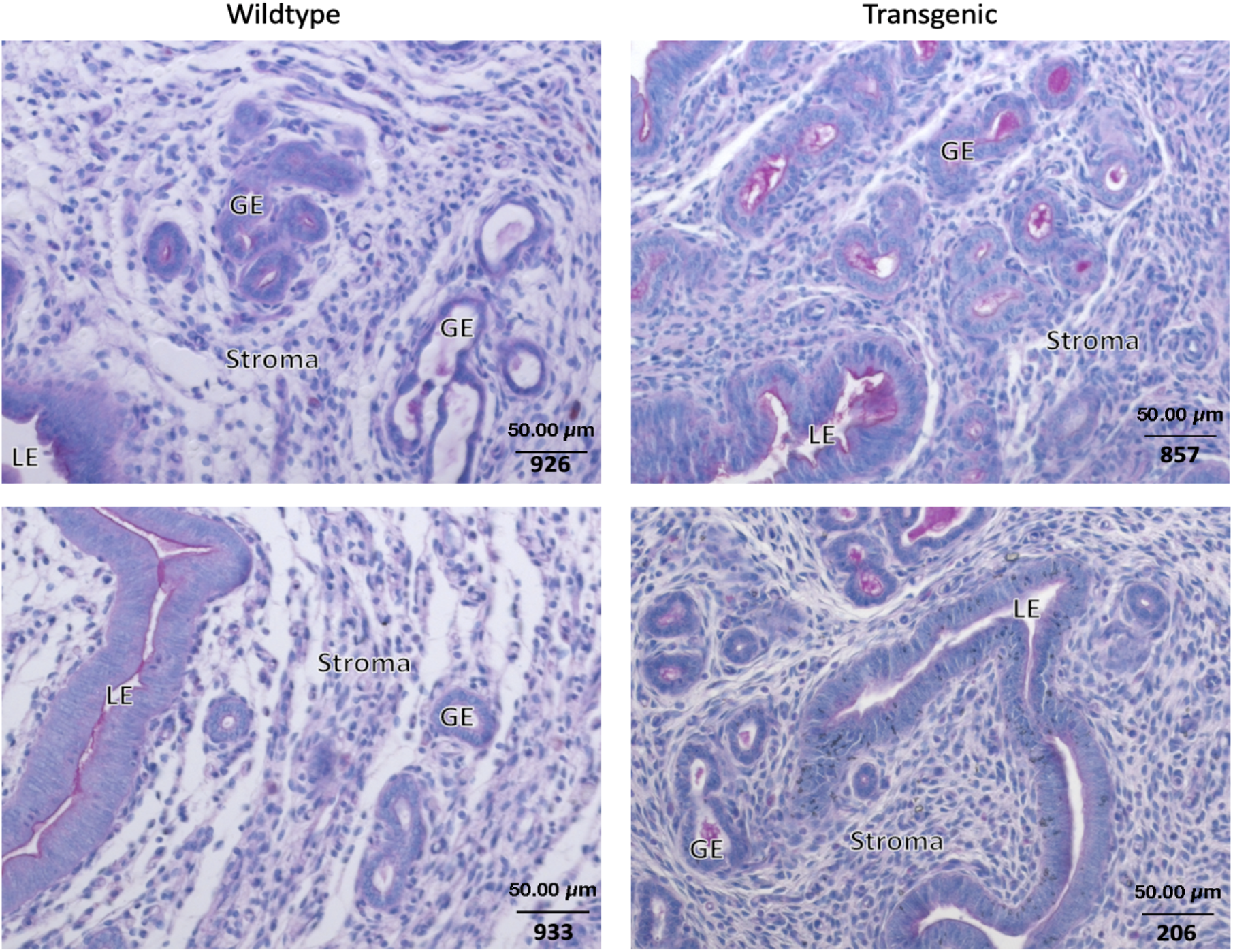
Transgenic line shows higher content of polysaccharides/mucins in proestrus uterine tissue. Periodic acid - Schiff stain of proestrus uterus from wildtype and transgenic lines detects a qualitative increase in the concentration of polysaccharide and mucin contents of the secretions in the lumen, particularly of that associated with the glandular epithelium. Magnification 20x.

### Wnt4 signals via non-canonical pathway in the proestrus endometrium

Given the *Wnt4* upregulation, and the observed transcriptional sequences, we asked what Wnt signaling pathway is used. The most prominent Wnt signaling pathway is the canonical pathway involving inhibition of ß-catenin degradation and promoting its translocation into the nucleus, where it acts as transcription factor of several Wnt target genes^30^. Wnt4 is known to signal via canonical as well as several non-canonical pathways in different contexts. We used immunohistochemistry in uterine tissue to examine whether the upregulation of *Wnt4* expression coincides with changes in nuclear localization of ß-catenin. We detected no nuclear localization of ß-catenin in the uteri of transgenic or wildtype mice during proestrus or estrus (Fig. S7A & B). Consistent with this result, there is no upregulation of the enhancer factor *Lef1* or member of the ß-catenin dissociation complex *Axin2* which are common targets of ß-catenin signaling^54^ (Fig. S7C), concluding that *Wnt4* likely causes described changes via one of the non-canonical pathways. An upregulation of *CamkIIb* (Fig. S7D) suggests the Ca2+/CamkII pathway may be involved, but the potential downstream mechanisms of Wnt4 activation require further attention.

### The uteri of genome edited mice manifest increased size during estrus

*Wnt4* mRNA is expressed at comparable levels in both genotypes during estrus (Fig. 2). However, we do note 41% increase in the uterine cross sectional area in the transgenic animals in estrus phase (P=0.05, N= (5 TG, 4 WT), t-test with heterogeneous variance; Fig. 7). As *Wnt4* expression is upregulated in transgenic animals in proestrus, we hypothesize that the size change observed in estrus is a downstream consequence of this upregulation. In the face of downregulated luminal epithelial proliferation, this volume increase is likely attributable to increased glandular activity or oedema.

**Figure 7.**
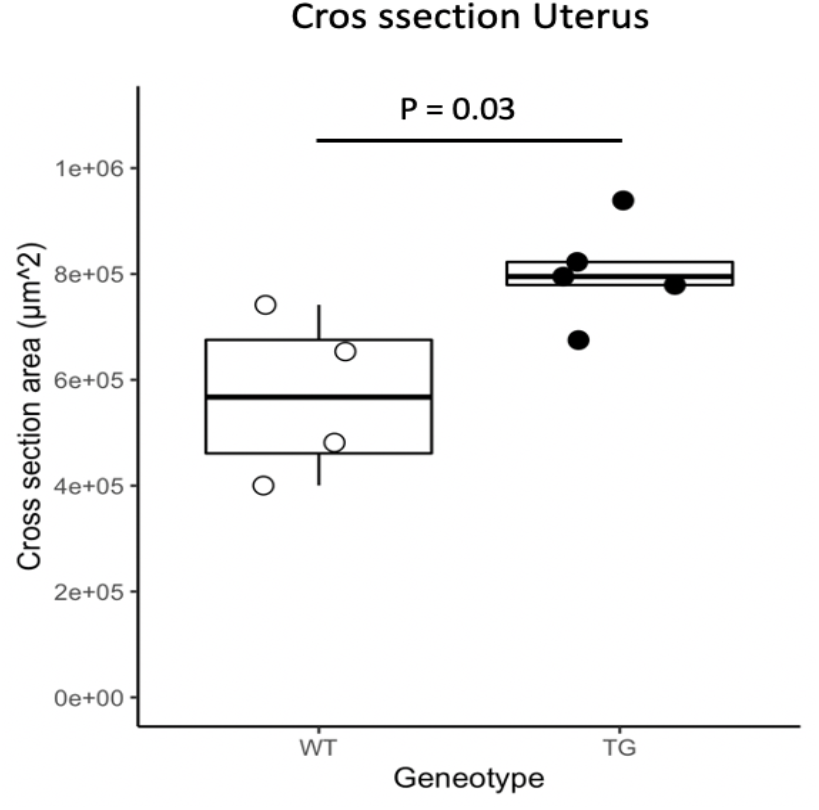
Increase in transgenic uterine size in estrus. Transgenic line showed an enlarged cross-sectional area of the uterine horn in estrus. As gene expression patterns indicated decreased proliferation, particularly in the luminal epithelium, differences in area likely correspond to increased secretion and luminal area along with possible oedema.

Overall, the observed proestrus expression profile in transgenic mice suggests that the activation of *Wnt4* by estrogen receptor in alternate allele genotype induces gene expression changes highly similar in spectrum to those mediated by progesterone in early pregnancy in mice and human, or in human secretory phase. In particular, we observe the downregulation of epithelial proliferation and upregulation of genes associated with stromal processes: decidualization, adhesion, extracellular matrix modification, invasibility and vascularization, as well as glandular activation. While several genes characteristic for decidualization are upregulated, the pattern doesn’t correspond to full decidualization. These changes precipitate the overall uterine volume increase in transgenic animals.

## DISCUSSION

In this study we report a functional analysis of the SNP rs3820282 that has been associated with multiple disorders of female reproduction and reproductive tissues. We have previously shown that the human alternate allele generates a high affinity binding site for the estrogen receptor alpha^19^. The present study aimed to uncover the immediate consequences of this allele substitution. Having created a transgenic mouse line with the human alternate allele, we took advantage of the natural estrogen dynamics during the mouse ovarian cycle to unravel the spatio-temporal context of mutational effects, and their direct consequences on gene expression. We found that the mutation causes upregulation of *Wnt4* in endometrial stromal cells, coinciding with the late proestrus estrogen peak. Importantly, the results demonstrate that the mutation activates uterine *Wnt4* expression in response to estrogen, independently of the presence of conceptus. The numerous changes in the downstream uterine gene expression suggest effects on two general processes: the suppression of proliferation in epithelium, and the upregulation of pathways involved with embryo receptivity and the uterine susceptibility to invasion.

### Transgenic gene expression changes and uterine biology

The mammalian uterus undergoes remarkable changes throughout the pregnant as well as nonpregnant cycle, mediated by the fluctuations in the ovarian steroid hormones estrogen and progesterone, and their respective receptors^55,56^. The transition from proestrus to estrus for example, is characterized by changes in transcription of a large number of genes, precipitating changes in morphology, tissue remodeling, adhesion and proliferation^57^. Stroma and epithelium react interdependently to changing hormone levels, as had been shown by the requirement of stromal Esr1 for epithelial estrogen action^58^.

Uterine *Wnt4* expression is estrogen-responsive in wild type mice^59^ and correspondingly, there are several Esr1 binding sites in the Wnt4 promoter. The single nucleotide polymorphism rs3820282 investigated here further modulates estrogen responsiveness of *Wnt4* in uterine stroma, specifically at the preovulatory estrogen peak in proestrus, thereby enhancing a non-canonical pathway in its target cells. Wnt4 subsequently acts as a downstream effector of progesterone signaling and is thus responsible for some of the effects otherwise associated with progesterone upregulation^60^. Specifically, Franco and coauthors^60^ showed that conditional ablation of Wnt4 increased apoptosis during stromal cell decidualization in mice and significantly reduced embryo implantation. Importantly, they found an effect of *Wnt4* upregulation on progesterone-regulated genes without upregulation of progesterone^56^ receptor, suggesting that Wnt4 mediates the effect of progesterone, consistent with the present results, with the exception that here Wnt4 is regulated in response to estrogen.

Implantation is a critical event, hence in species with spontaneous ovarian cyclicity several recurring uterine changes have evolved in preparation for implantation. In most mammals, some uterine changes precede the presence of the conceptus and thus occur also in the nonpregnant cycle (e.g., edema), whereas others require copulation or the presence of conceptus. In humans, the preparatory changes are more extensive and include stromal decidualization, which in almost all non-primate mammals requires the presence of conceptus. Mammalian preimplantation uterine changes include actin cytoskeleton remodeling and reorganization of focal adhesions and junctional complexes in stromal cells, loss of epithelial polarity, changes in apical membrane and increase of glycogen production in luminal epithelium and stroma, as well as changes in vascular permeability and angiogenesis^61–64^. As several of these processes are mediated by progesterone, the observation of enhanced pro-implantation signature downstream of *Wnt4* expression is plausible. A surprising aspect is that these progesterone-induction-like changes precede the progesterone-dominated luteal phase. In mice, edema and angiogenesis increase during estrus (coinciding with increase in *Wnt4* expression), while further uterine remodeling occurs in days following copulation and fertilization, as a functional corpus luteum starts secreting progesterone. The upregulation pattern in a proestrus transgenic mouse therefore suggests an advanced induction of the preimplantation processes, which in absence of the conceptus, is broadly reminiscent of the human condition of spontaneous decidualization in humans. The early expression of *Wnt4* may enhance the receptivity by increasing the *invasion susceptibility, i.e., invasibility* of the uterus for implantation, thus explaining the positive effect of this mutation on human pregnancy outcomes^19^.

### Comparability to human

Predicting the comparability between model organisms and humans requires understanding of the shared, and accounting for the species-specific mechanisms. The mouse and human ovarian cycle and pregnancy have diverged during evolution. While we do not have a full mechanistic understanding of these differences, we know that many aspects of receptivity are conserved across mammals^61^. This is due to a shared evolutionary origin of invasive implantation in the stem lineage of placental mammals^65–67^, despite the likely independent origins of mechanisms controlling subsequent gestation length^67^. The presented consequences of *Wnt4* upregulation are driven by the dynamics of ovarian preovulatory estrogen peak, which is shared between mouse and human. This speaks for the relevance of the present findings for the human condition. Understanding the downstream effects of a SNP in the mouse uterus may therefore enable prediction of the spatio-temporal contexts in which this SNP exerts its effects on human pregnancy and physiology.

### Relation to endometriosis and reproductive cancers

Our findings provide the most direct mechanistic evidence to date for a link between this particular SNP and reproductive traits. Our results imply a two-pronged effect of the mutation: suppression of proliferation in the epithelium and enhancement of pro-implantation/invasion processes in the stroma. The association of the SNP with cancer and endometriosis may involve one or both of these. We suggest that the reported correlated associations of the mutation with reproductive cancers and endometriosis may be mediated through the second prong: the enhancement of pro-implantation processes, which make stroma more permissive to invasion by embryo but also by cancer and ectopic endometriotic tissue. This suggestion is inspired by the correlation across species between endometrial permissiveness to embryo implantation and vulnerability of peripheral stroma to cancer invasion, where species with more invasive placentation are more prone to invasion by metastases^68–71^. Here we hypothesize that this relationship is not only a species-level phenomenon but applies within human populations; individuals with increased uterine embryo invasibility may be more prone to cancer metastasis and endometriosis.

Supporting the increased likelihood of cancer invasion in the transgenic line, is the set of the stromal dysregulated genes which match the expression pattern associated in the literature with stromal support of cancer invasion. The processes involved are matrix remodeling by proteases and cell-matrix adhesion (increased integrins and periostin), vaso-activation (increased *Fst*, *Rgs^72^*, *Serpine1^52^*), energy provisioning (increased *Slc2a12*^74^), and promotion of invasion (decreased *Spink12^73^*, *Pdgfra^74^*). Among the known pathways that influence tissue invasibility is also Wnt signaling^75^. Thus, differences in tumor microenvironmental expression, not only in the cancer cells themselves, may explain the association with cancer invasion. Awareness of the importance for cancer invasion of supportive stromal microenvironment has increased in recent research^76–78^.

While the dysregulation of *Wnt4* is temporally restricted by the estrogen peak, it is not necessarily spatially restricted to the uterus. It is plausible that the underlying processes are shared with other tissues expressing ESR1 receptors, even if the downstream effector genes may differ. The periodically elevated estrogen in the estrous cycle could thereby affect either the cancer cells themselves or the stroma exposed to and interacting with cancer cells. Such menstrual phase-dependent cancer invasibility has previously been suggested with respect to the propensity towards post-resection metastasis in breast cancer^79^. These effects have been suggested to be due to cycle-phase-dependent fluctuation in angiogenic factors (e.g., VEGF) and factors promoting epithelial-to-mesenchymal transition, downstream of estrogen^80^. The results of the present study suggest that vulnerability to this effect may be genetically enhanced by the alternate allele at the locus rs3820282.

Estrogen responsiveness and *Wnt4* activation are common characteristics of many cancers, even if the mechanisms downstream of Wnt4 are variable^81–84^. For example, a recent study has described the co-option of WNT4 under the control of ESR1 in the invasive lobular breast carcinoma, with downstream positive effects on tumor cell proliferation, mTOR signaling and mitochondrial function^17,85^ – the effects we do not observe in the transgenic mouse uterus, which may be due to the absence of cancer cells.

The epithelial prong of proliferation suppression in our mouse model contradicts the expectation from endometriosis. Endometriosis has been associated with the failure of progesterone receptor to induce genes counteracting estrogen-driven cell proliferation in the secretory phase, i.e., the progesterone resistance^26^. The strongest dysregulation of uterine gene expression in endometriosis was detected in the early secretory phase, whereby a significant *Wnt4* dysregulation is rarely reported (but see Liang et al.^89^, for *Wnt4* decrease). Interestingly, endometriosis-associated gene dysregulation in early secretory phase matches dysregulation in the transgenic mouse endometrium in the present study in terms of affected gene-set, above all the progesterone-regulated genes (e.g., *Errfi1, Bcl6, Cited2, Irs2, Tgfb2, Gpx3, Tk1^26^*). However, these genes are dysregulated in opposite directions. This is an intriguing result, as the support for the causal role of the locus rs3820282 in endometriosis is strong^16,27^. Several factors could explain the mismatch between the effect in mouse proestrus, and its predicted effect on endometriosis: the species context (human vs. mouse), or the differing proportions of stromal and epithelial cells between mouse endometrium and endometriotic lesions. Alternatively, the associations with endometriosis and cancer may primarily stem from the effects of the SNP other than those on epithelial proliferation such as enhancement of pro-implantation/invasion- a hypothesis we propose.

Wnt4 has previously been proposed as the plausible causal candidate for endometriosis because of its genomic proximity to rs3820282 and its role in the development of the female genital tract^86^, which could precipitate a tendency towards reproductive defects^22,87^. However, functional genetic follow-up on rs3820282 in the context of endometriosis did not confirm this effect. Luong and colleagues^27^ and Powell et al.^25^ tested the genes flanking rs3820282, and found that the alternate allele was associated with downregulation of LINC00339 (RNA gene absent from the mouse locus) in whole blood and endometrium, and upregulation of *CDC42* in whole blood. WNT4 was not detected in whole blood samples (Brisbane genetics study^88^) and also no effect of the rs3820282 genotype on *WNT4* expression in endometrium was detected. Chromosome conformation capture (3C) in Ishikawa endometrial cell line found looping of the regulatory region with the SNP to *CDC42* and LINC00339^25^. There are several plausible sources for the discrepancy between these results and our current findings. First, while authors accounted for sample variation due to different stages of the menstrual cycle, only 16 of 132 samples were collected in the early secretory phase, somewhat close to the stage in which we show the effect. Second, 3C approach would not reveal the regulation of WNT4 as the SNP is in the *WNT4* intron itself. Finally, the Ishikawa cell line is derived from adenocarcinoma and thus has an uncertain cellular identity. These cells have been reported not to express *WNT4^89^* – rendering this line unsimilar to endometrial cells. Thus, the role of *WNT4* as a mediator of the locus’ effect could not be excluded by this study.

### Conclusion

We suggest that antagonistic pleiotropy associated with *rs3820282* derives from a common mechanistic effect, namely an increased tissue support for invasion, occurring in different contexts. In the context of embryo implantation, the mutation enhances a normal process. In the context of reproductive cancers and endometriosis, it may not directly affect the origin or proliferation of cancer cells, but it does decrease the host tissue’s ability to resist the invasion, or even provide a supportive context. The idea of an evolutionary trade-off between the mammalian invasive implantation and the vulnerability to cancer metastasis is gaining support in recent research^69^. What may be common to both processes is the shared active role of the stromal microenvironment in regulation of invasion, either the embryo or cancer cells. The current study adds to the mechanistic understanding of this phenomenon by pointing out that such permissiveness to cancer may not be persistent but is limited to specific phases in the ovarian cycle, which modify not only the uterine, but potentially multiple estrogen responsive tissues. This differs from the idea of pre-metastatic niches in that the permissiveness of tissues to cancer invasion is not induced by the cancer cells themselves but is rather a side-effect of evolved uterine permissiveness and thus occurs, without consequences, also in the absence of cancer (or embryo). This enables a potentially different approach to restricting metastasis of estrogen-responsive cancers - by addressing its spatio-temporally local active support rather than modifying a ubiquitous growth process.

## METHODS

### Animal husbandry

Animals were kept under 12:12 light:dark cycle and bred within genotype in house. Offspring were weaned at three weeks and reared separately by sex, housed no more than four animals per cage and fed standard mouse chow diet ad libitum. Animals were euthanized by asphyxiation (CCHMC) and cervical dislocation (UoV), consistent with animal care protocols at CCHMC and the UoV, immediately prior to tissue harvest. All procedures were approved by the relevant animal care and protection boards (CCHMC: IACUC2016-0053; UoV: Austrian Ministry for Science and Education).

### Generating transgenic mice

The aim of gene editing was to replace the mouse allele at the location in the mouse genome corresponding to rs3820282 in humans, with the human allele. The position rs3820282 and its flanking sequences are 98% conserved between the human and mouse genome (Fig. 1). For gene editing, we selected the sgRNAs that bracket the corresponding site in mice based on the on- and off-target scores from the web tool CRISPOR^90^. The selected sgRNA is constructed in a modified pX458 vector^91,92^ that carries an optimized sgRNA scaffold^93^ and a high-fidelity Cas9 (eSpCas9 1.1)-2A-GFP expression cassette^94^. Individual sgRNA editing activity is validated in mouse mK4 cells, using a T7E1 assay (NEB), and compared to the activity of Tet2 sgRNA that has been shown to modify the mouse genome efficiently^95^. Validated sgRNAs are in vitro synthesized using MEGAshorscript T7 kit (Life Technologies) as previously described^96^. Injection was made into the cytoplasm of one-cell-stage embryos of the C57BL/6 genetic background using the method described previously^96^. Injected embryos are immediately transferred into the oviductal ampulla of pseudopregnant CD-1 females. Live born pups are genotyped by PCR and then further confirmed by Sanger sequencing.

### Genotyping

DNA for genotyping was isolated from tail clips and a PCR was run using the following primers: fwd: GCCTCAGAGGAATTGCGAGC and rev: GGATAGCCAACAGTGTAGCTGG. We used a high-fidelity polymerase Regular Phusion (NEB (M0530)) with high annealing temperature, and GC buffer. PCR cycling was as follows:

98C hold until load samples-

Hot Start

1. 98°C 2’
2. 98°C 15”
3. 65°C 20”
4. 72°C 30”
5. 30 cycles steps 2-4
6. 72°C 5’
7. 15°C until emptied

The products were digested with restriction enzyme Tsp 451 (NEB R0583), which cuts if the alternate T allele is present or with Msc1 (NEB R0534), which cuts if the C allele is present, Only clean bands were purified with QIAquick PCR purification kit (Qiagen) and sent for Sanger sequencing.

### Gestation phenotype

To assess the effect on gestational length, the pregnancies of homozygous females bred to the males of the same genotype were observed. The midnight preceding the morning of plug detection was counted as beginning of pregnancy, and the females were monitored to determine the start of parturition as appearance of the first pup. The litter size was assessed after the labor ceased and checked on the next day.

### Tissue Collection

Animals were bred and tissue collected and harvested at two institutions (University of Vienna, UoV, and Cincinnati Children’s Hospital Medical Center, CCHMC). Analyses were repeated with samples from each institution independently, and the results have been consistent.

#### Proestrus and estrus

Adult non-mated female mice (aged 2-8 months) were monitored by daily vaginal swabs to determine the stage of estrous cycle^97^. Animals of each genotype at the locus of interest were harvested in proestrus and estrus, respectively. Uterine horns and ovaries were collected from each individual, one of each was placed in 4% PFA for histological and immunohistochemical assessment, the second was flash-frozen and/or stored in RNAlater for RNA extraction.

#### Decidualization

To assess gene expression during decidualization in vivo, we mated females of known genotype (TG or WT) with males of the same genotype and separated them upon detecting the copulatory plug in the morning examination, counting noon of the detection day as 0.5 day post copulation (dpc). We harvested only the pregnant uteri at 7.5 dpc and preserved the tissue by flash freezing in liquid nitrogen for later RNA extraction and qPCR.

#### Primary cell isolation and culture

We isolated primary mouse uterine stromal fibroblast cells using a standard protocol^98^, with the difference that uteri were harvested from mice in proestrus, as established by daily vaginal swabs^97^. In brief, uterine horns were harvested and digested with pancreatin and trypsin. The detached epithelial sheets were removed from the solution and the remaining tissue underwent two consecutive rounds of digestion in collagenase (30 min each), washing and filtering of the cell-containing media through 40 μm nylon mesh, spinning and seeding in ESF medium. Mouse ESF culture growth medium consists of equal parts of Dulbecco’s Modified Eagle Medium and Nutrient Mixture F12 (DMEM/F12) with phenol red, L-glutamine and 4-(2-Hydroxyethyl) piperazine-1-ethanesulfonic acid, N-(2-Hydroxyethyl)piperazine-N′-(2-ethanesulfonic acid) (HEPES), 10% FBS, 0.5 μg/mL amphotericin B, and 100 μg/mL gentamicin. Cells were grown to confluence in 6-well plates. Before the extraction, the growth medium was removed, and the cells were rinsed with PBS.

### Histology

#### Sample preparation

Fresh tissues were fixed in 4% PFA overnight at room temperature. They were then washed with PBS and transferred to gradually increasing concentration of ethanol (30-50-70%). Finally, tissues were then processed and embedded in paraffin blocks. For histological examination, slides were cut at 5 μm thickness and stained by hematoxylin-eosin staining using standard protocols.

#### Immunohistochemistry

Paraffin slides were incubated at 60°C overnight, washed in 3 changes of Xylene to remove paraffin, and rehydrated in gradually decreased Ethanol (100-95-70%) and PBS. Antigen retrieval was conducted using citrate buffer (pH 6.0), followed by H_2_O_2_ to remove endogenous peroxidase activity, and blocking serum to avoid non-specific binding. In all cases, non-conjugated primary antibody was made visible with a secondary antibody with fluorescent fluorophore. The following rabbit-raised anti-mouse primary antibodies were used. Anti-Mki67(Abcam; ab16667), concentration 5μg/ml and anti-beta catenin (Abcam; ab16051), concentration 1μg/ml. In both cases we used a goat-raised, anti-rabbit, fluorophore-conjugated secondary antibody (Abcam; ab150077), concentration of 2 μg/ml.

#### RNA scope (*in situ*)

Commercial RNA scope kit for mouse *Wnt4* RNA was purchased from Advanced Cell Diagnostics, Inc. (ACD). In situ hybridization was performed on paraffin-embedded tissue, following the manufacturer’s instructions.

### Profiling gene expression

#### RNA isolation

From frozen uterine horn or ovary approximately 20 mg of tissue was lysed with Precellys Evolution homogenizer at 6500 rpm 2 × 20s with a stainless-steel bead and 0.04M ditiotheritol. RNA was isolated using the RNeasy mini kit (Qiagen) according to the manufacturer’s instructions. RNA was subsequently stored at −80°C. From confluent primary cells, RNA was isolated using the mirVANA kit (Thermo Fisher Scientific), according to the manufacturer’s instructions. RNA was subsequently stored at −80°C.

#### qPCR

RNA expression profiles across estrous cycle of genes adjacent to the SNP (Cdc42 and Wnt4) were established by qPCR and compared between the genotypes. RNA was treated with TURBO DNase (Thermo Fisher) to remove any genomic DNA and converted to cDNA with High-Capacity cDNA reverse transcription kit (Thermo Fisher) following standard protocols. qPCR was performed with TaqMan primer probes (Thermo Fisher) with VIC dye for the *GapDH* internal control and multiplexed with FAM for *Wnt4*. Proestrus and Estrus samples were run on an AriaMx qPCR with Brilliant III Ultra-Fast master mix (Agilent). 7.5dpc and decidual cell lines were analyzed with TaqMan Gene Expression master mix (Thermo Fisher) on a Mastercycler realplex (Eppendorf). Relative gene expression (2^−ΔCt^) was calculated with the respective associated qPCR analysis software. Statistical analyses and visualization of qPCR results were done using R version 4.1.1^99^. Statistical significance was evaluated by Wilcoxon-Mann-Whitney rank sum test. Only samples with values differing by more than 10 SD were removed.

#### Transcriptome analysis (RNA Seq)

RNA from four TG and three WT proestrus uteri was collected as described above. Library preparation and sequencing was conducted by the CCHMC sequencing facility on an Illumina NovaSeq 6000 to the depth of 30 million paired reads of 100bp length. Reads were aligned to the mouse GRCm38 genome^100^ with STAR using settings -- outFilterMultimapNmax 1 -- quantMode GeneCounts^101^. Gene expression was analyzed with DEseq2^102^ and an adjusted p value >0.05 was used to identify differentially expressed genes, shrunken log2FC values were used to account for lowly expressed genes^103^ (Table S1). Relationships among samples were investigated with variance stabilizing transformed data and batch effects removed with limma^32^ using a principal components analysis as well as hierarchical clustering of Pearson correlation between samples. Gene set enrichment^104^ was analyzed with clusterProfiler^105^ using the Hallmark collection from the Molecular Signature Database^32^, using ranked shrunken log2FC (Table S2). Data was analyzed with R version 4.1.1^99^. All raw data has been deposited to European Genome-Phenome Archive (available at publication).

## Acknowledgement

The work was funded by the March of Dimes Ohio Collaborative Project to L. Muglia (#22-FY14-470) and Austrian Science Fund to MP (FWF #P33540). We particularly acknowledge the help in histology by Gale Macke of Cincinnati Children’s Hospital and the research support by Elisabeth Rauscher at the University of Vienna.

## Contributions

MP, GPW, LM developed the idea and planed the research, LM, AMZ, CEM-G and MP performed the experiments and the analyses, Y-CH helped with generating transgenic lines, NM and GD helped with the genotyping, LM, GD, FK and JM were instrumental in maintaining the colony and collecting the tissue, NZ helped with organizing the samples and validation, DS contributed to localization of the mutational effect, GZ crucially contributed to interpretation of genomic signal, MP and CEM-G wrote the draft. All authors have reviewed and discussed results, interpretation and formulation.

## Supplemental Figures

**Figure S1.**
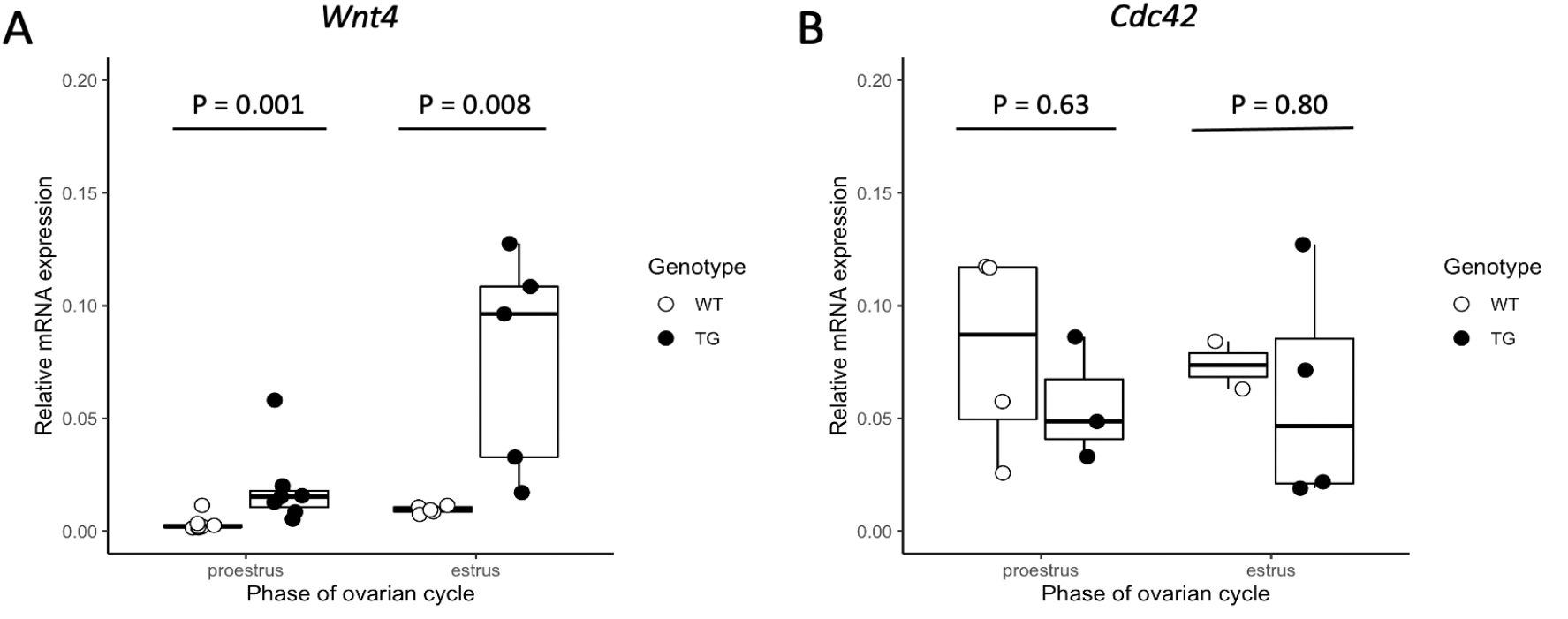
SNP influences expression of Wnt4 in the uterus in the replicate transgenic line. In the additional mouse line generated, the transgenic allele resulted in similar mRNA gene expression changes measured by qPCR compared to the primary line used in experiments (Fig. 2). A) Significant increase in *Wnt4* expression changes were also found in the replicate transgenic line (KI-2). This difference in expression persisted into the estrus stage in this line. B) As in the other transgenic line, there was no significant difference in *Cdc42* expression. As this line did not breed as robustly it was not used for subsequent experiments. Nevertheless, this demonstrates the reproducible effects of the SNP on *Wnt4* expression.

**Figure S2.**
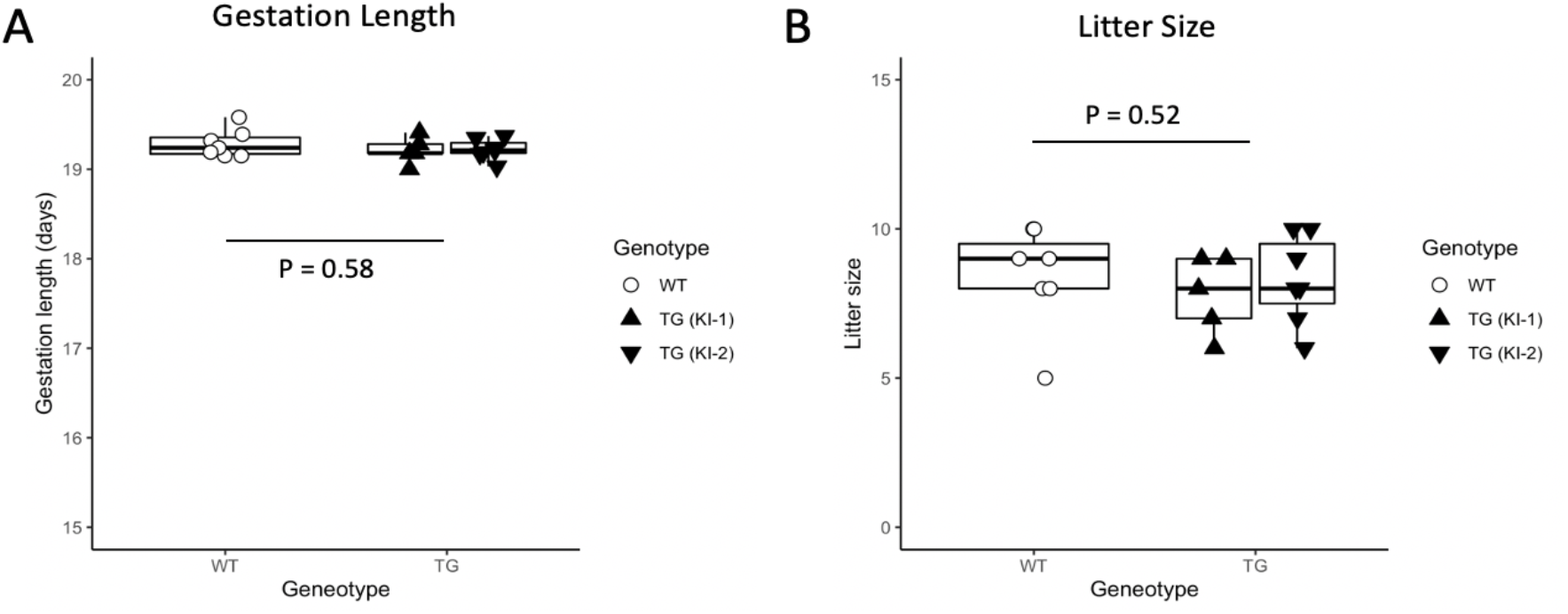
No differences in gestation length or litter size in transgenic lines. A) Gestation length and B) litter size was similar across all genotypes.

**Figure S3.**
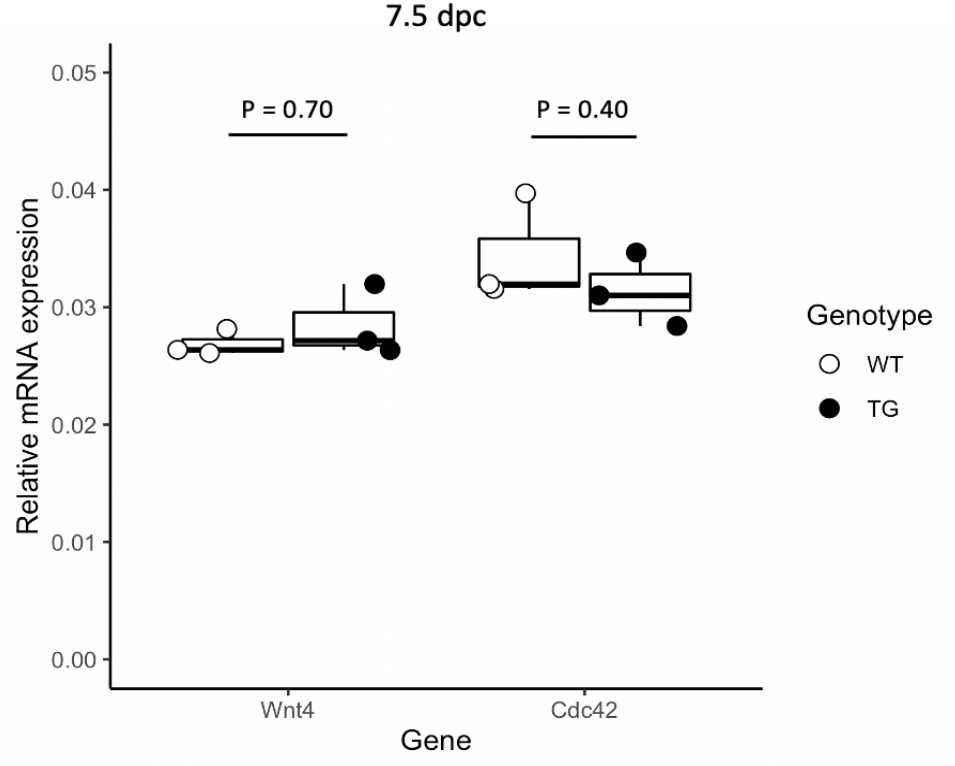
No difference in Wnt4 expression in the transgenic 7.5 dpc uterus. mRNA expression, measured by qPCR in bulk uterine tissue from 7.5 dpc pregnant animals, does not indicate significant upregulation of *Wnt4* in transgenic animals.

**Figure S4.**
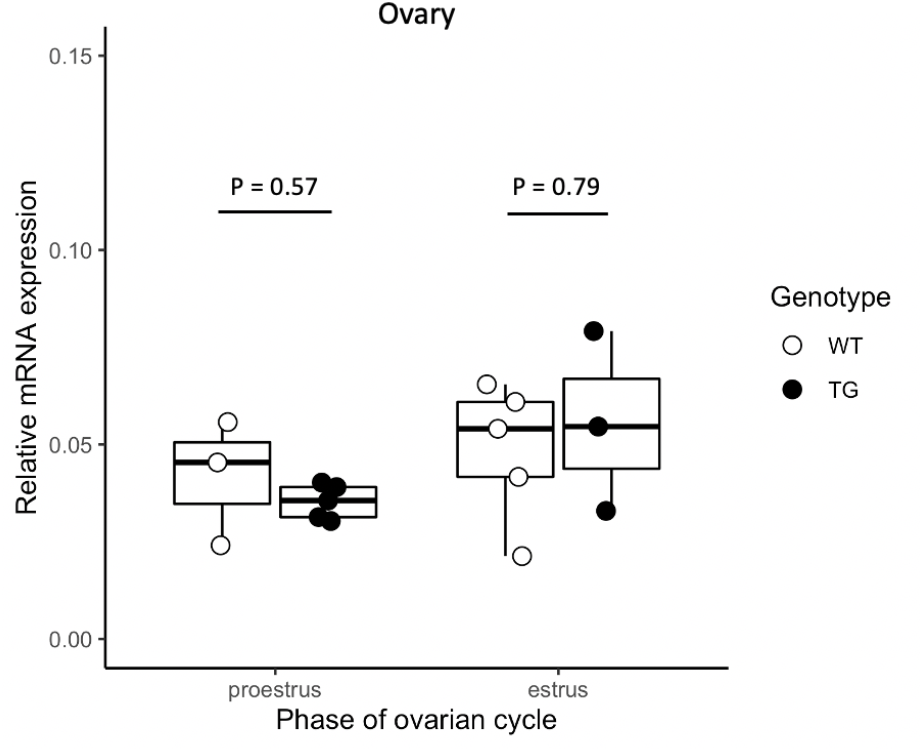
SNP does not influence expression of Wnt4 in the proestrus and estrus ovary. mRNA expression, measured by qPCR in bulk ovarian tissue from estrus and proestrus, does not indicate significant upregulation of *Wnt4* expression in the ovary of transgenic animals.

**Figure S5.**
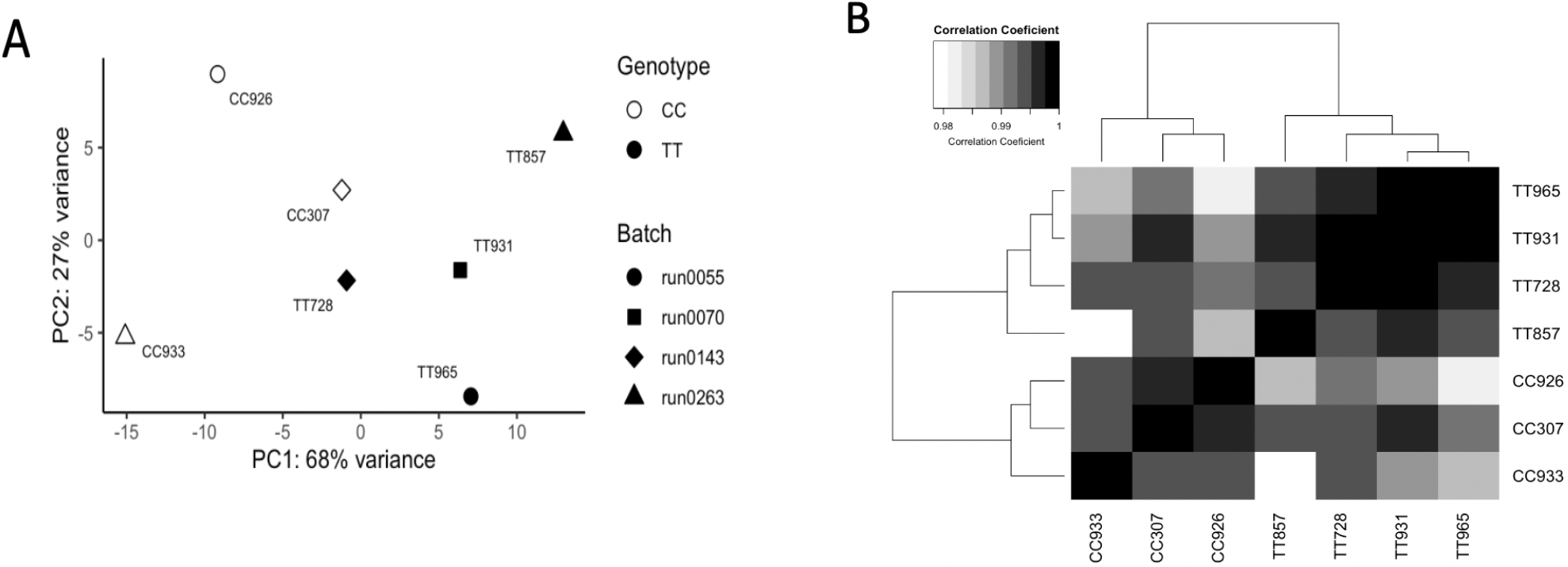
Reproducibility of transcriptome. SNP transgenic genotype corresponds to gene expression differences among samples following batch correction and normalization. A) The first principal component explained 68% variation and generally separated samples based on genotype in batch corrected variance stabilized gene counts. B) Pearson correlation between all samples was greater than 95% with hierarchical clustering distinguishing samples based on genotype.

**Fig S6.**
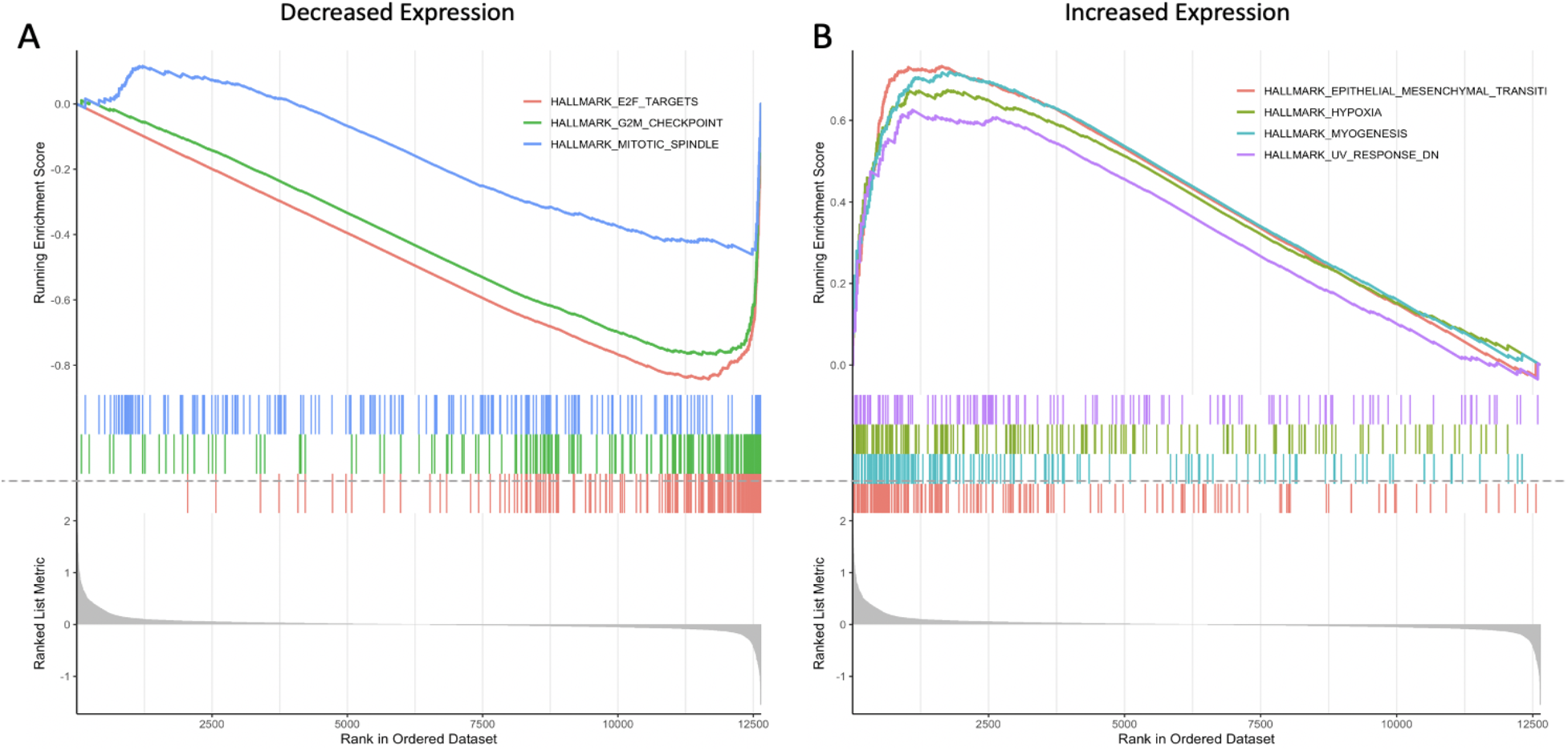
Distinct patterns of functional enrichment for differentially expressed genes. A) Genes with decreased expression in the transgenic line (i.e., higher expression in the wildtype) were enriched for functions associated with mitosis and cell cycle progression including E2F targets, G2M checkpoint, and Mitotic spindle. B) Genes with significantly increased expression in the transgenic line were enriched for a variety of functions including epithelial mesenchymal transition, hypoxia, myogenesis, UV response down (DN). All significantly enriched gene sets can be found in Table S2.

**Figure S7.**
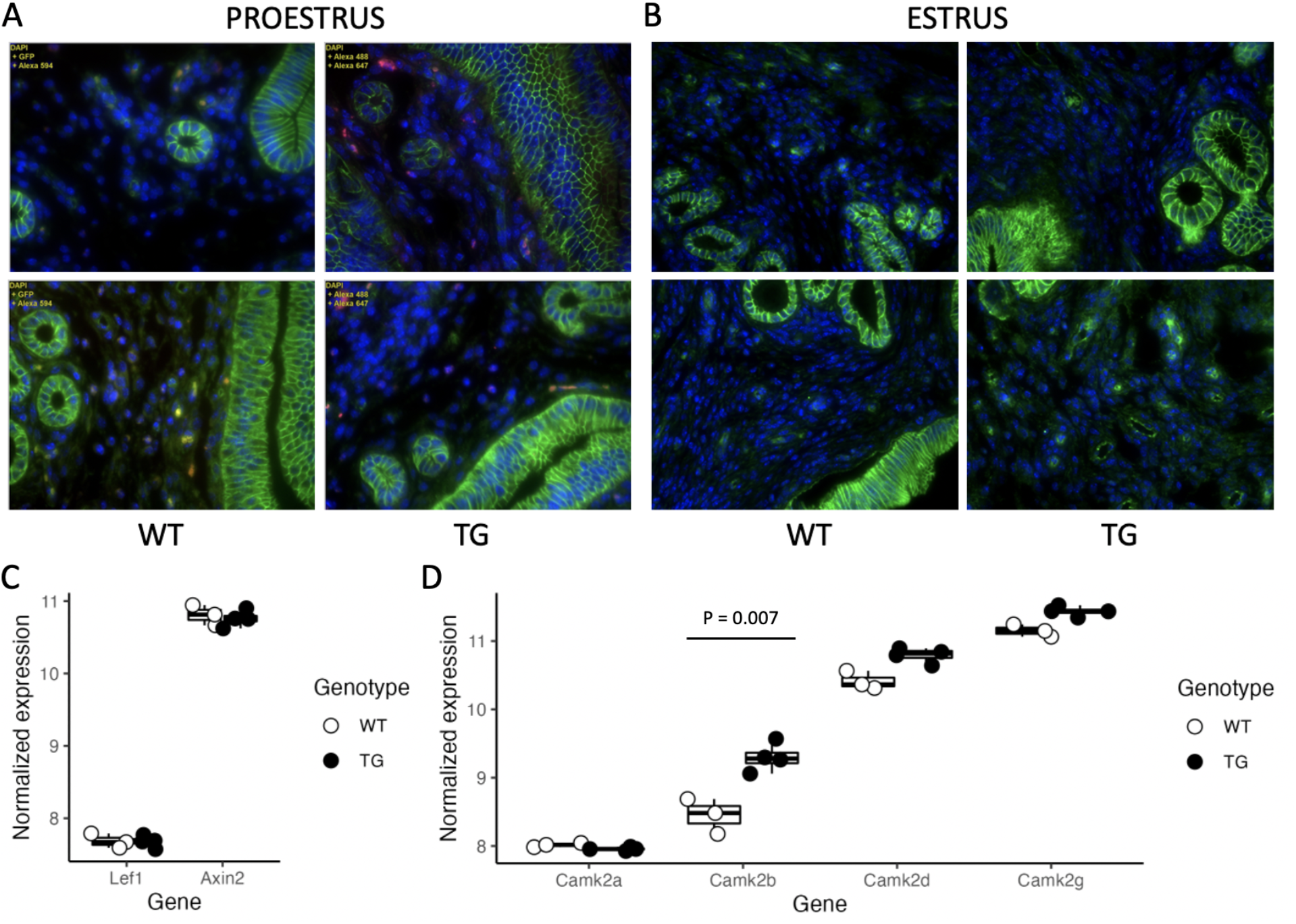
Support for non-canonical Wnt4 signaling. (A) Immunohistochemistry using ß-catenin antibody shows no translocation of ß-catenin into the nucleus in wild type or transgenics, in proestrus or estrus. Lack of ß-catenin translocation indicates that the canonical Wnt4 pathway is not activated. (B) Supporting the absence of canonical Wnt4 pathways, enhancer factor *Lef1* expression is at the limits of detection with insufficient expression to evaluate expression changes and high expression of *Axin2*, a member of the ß-catenin dissociation complex, with no evidence of increased expression associated with ß-catenin activation. (C) One of the non-canonical signaling pathways is via increased intracellular calcium which can result in phosphorylation of CamkII. Genes for the four alternative CamkII subunits were detected, with increased expression of *CamkIIb* in the transgenic line.

## Supplemental data

**Table S1 RNAseq Data** (Analyzed data) Batch corrected and normalized expression values along with DEseq2 analysis of differential expression. (Filtered raw data) Count data filtered for genes expressed (at least 1 count) in each sample.

**Table S2 Hallmark gene set enrichment** All Hallmark gene set functional categories enriched within differentially expressed genes and associated statistical analyses.

